# Glioblastoma model mouse rescue from mortality by brain-targeted catalytically-enhanced angiotensin-converting enzyme 2 mutant-antibody fusion protein

**DOI:** 10.64898/2026.11.03.763994

**Authors:** Pete Heinzelman, George R Nageeb, John Choi, Justin Liu, Si Yeon Lee, Philip A Romero, Michael Lim

## Abstract

We have discovered that an engineered angiotensin-converting enzyme 2 (ACE2)-brain-targeted antibody fusion protein, referred to as ACE2dm-s1, improves survival of CT-2A xenograft glioblastoma (GBM) model mice when administered in combination with a small molecule STING agonist cancer therapeutic. Particularly striking about these survival results is that prior efforts that used losartan, a small molecule angiotensin receptor antagonist that acts on the same Renin-Angiotensin System (RAS) axis that is affected by the action of ACE2dm-s1, showed that this small molecule provided no survival benefit when used as an element of combination therapy in the same CT-2A GBM model used in our experiments. The improved CT-2A mouse survival, which was not observed for ACE2dm-s1 or STING agonist monotherapy, combines with our flow cytometry study results that revealed an ACE2dm-s1-mediated shift of brain immune cell populations toward an anti-tumoral composition to establish ACE2dm-s1’s utility as a tumor microenvironment (TME) modulator that could serve as an adjuvant in increasing the efficacy of wide range of cancer treatment modalities including chemotherapeutic agents, immune checkpoint inhibitors, and CAR-T cells.

## Main Document

We have uncovered a new approach to modulating the renin-angiotensin system (RAS) axis in cancer therapy that improves survival in the CT-2A xenograft mouse model of glioblastoma (GBM) and can also be broadly relevant in treating systemic solid tumor cancers. The majority of current efforts that target the RAS in cancer therapy, which aim to impact signaling pathways associated with cells residing within the tumor microenvironment (TME) to reverse the TME’s immunosuppressive nature, prevent or attenuate pathological fibrotic processes, and increase tumor blood perfusion through vasodilative processes (1), seek to elicit these effects by attenuating activation of the angiotensin II type1 receptor (AT1R) by the angiotensin II (Ang-II) peptide. Small molecule AT1R antagonists, such as the hypertension medication losartan, have been the most commonly used pharmacologic agents in both animal model experiments (2) and human trials, such as NCT01821729, in which the RAS has been pursued as a therapeutic node. RAS modulation via AT1R antagonism however, does not touch on all plausible angles for influencing the RAS axis.

Activation of the Mas receptor (MasR) by its cognate angiotensin-(1-7) (Ang-(1-7)) agonist peptide (3) can have salutary effects, e.g., attenuation of transforming growth factor beta1 (TGF-β1)-driven fibrosis, reduction of NF-κB-promoted production of inflammatory cytokines, and increased vasodilation via elevated endothelial cell production of nitric oxide, that counter undesired AT1R activation and rebalance the tumor local RAS (Fig 1a) to reduce tumors’ ability to, by way of upregulating genes such as *agt* that are involved in Ang-II biosynthesis, leverage increased local production of Ang-II in achieving AT1R hyperactivation that reduces tumor susceptibility to endogenous immune responses and pharmacologic interventions. Although activation of MasR via administration of exogenous Ang-(1-7) has been pursued in rodent cancer models and human clinical trials (4), efforts in this area have subsided over the past decade likely due to Ang-(1-7)’s nonideal pharmacokinetics, i.e., half-life in blood on the order of minutes, presenting a challenge to clinical translation of this peptide for cancer therapy.

**Fig 1.**
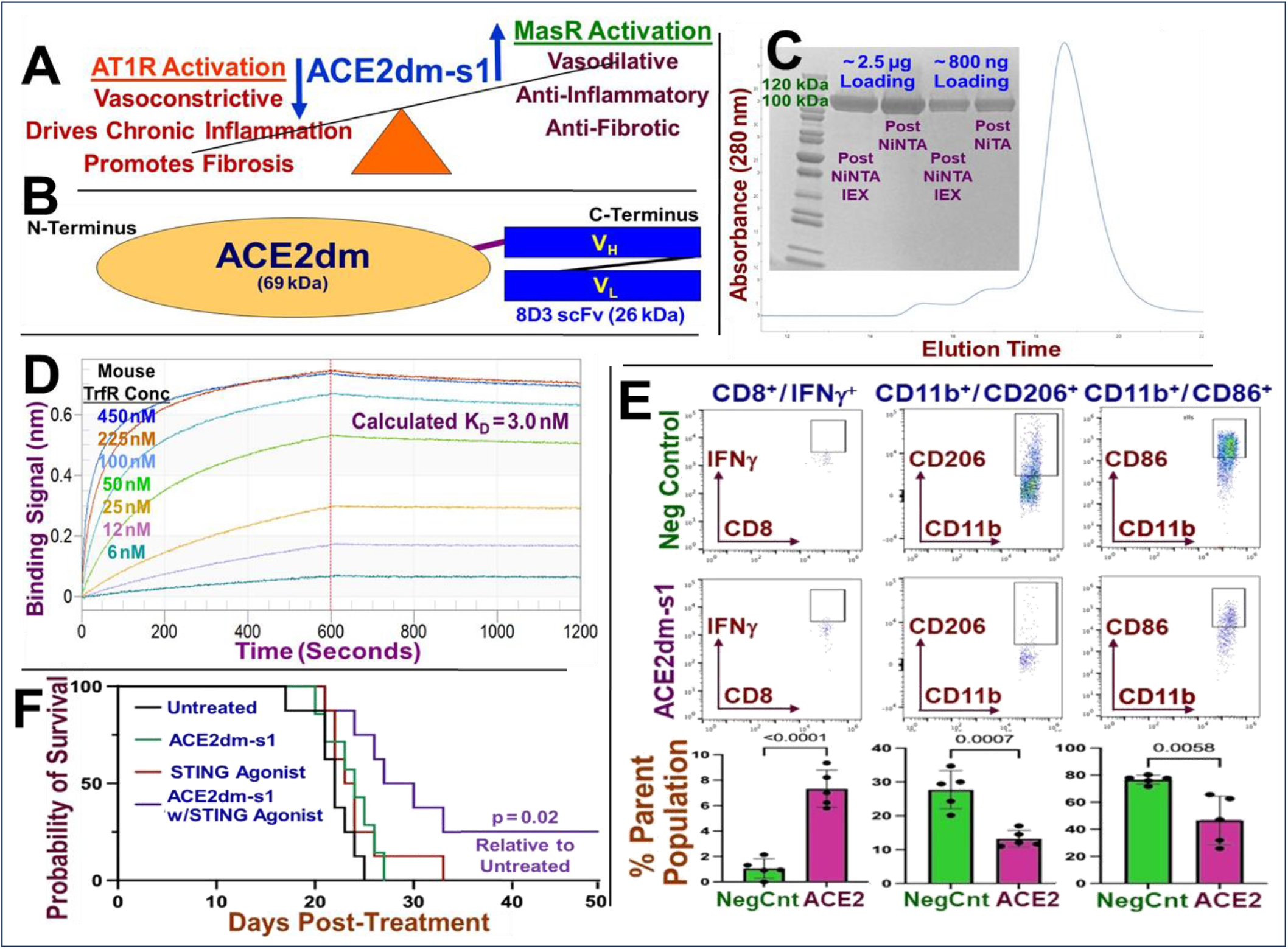
**(A)** MasR axis rebalancing by ACE2dm-s1. (**B**) ACE2dm-s1 schematic. (**C**) SDS-PAGE analysis for ACE2dm-s1 post one-stage (NiNTA) and two-stage (NiNTA and ion exchange (IEX)) purification and size exclusion chromatography trace for NiNTA/IEX-purified ACE2dm-s1. Bands appear near theoretical molecular weight of 95 kDa. (**D**) Biolayer interferometry response curves for immobilized ACE2dm-s1 and various concentrations of mouse TrfR free in solution. (**E**) Representative flow cytometry dot plots (1,500-15,000 events per parent population) and percent parent population (CD45^+^/CD11b^+^ for macrophages and CD3^+^/CD8^+^ for T cells) values for brain immune cells isolated from CT-2A mice. Error bars denote deviation for n=5, p-values obtained from Student’s t-test. (**F**) Survival curves for CT-2A mice receiving five-day monotherapy or ACE2dm-s1 (3.5 mg/kg - intraperitoneal) with 2’,3’-cGAMP STING agonist (3 µg per injection - intrathecal) combination therapy treatment course. n = 8 or n = 7 mice per group.

Due to its ability to hydrolyze the Ang-II octapeptide (DVIYIHPF) to produce Ang-(1-7) (DVIYIHP), the human angiotensin-converting enzyme 2 (ACE2) carboxypeptidase holds appeal as a RAS-modulating therapeutic that simultaneously reduces AT1R activation and increases activation of MasR and possess more desirable pharmacokinetic properties than Ang-(1-7). Wild type ACE2 however, hydrolyzes a range of peptide substrates, including apelin peptides important for cardiac function, with this off-target activity creating potential for side effects and making the enzyme susceptible to reduction of Ang-II hydrolysis rate by competition effects in the in vivo mixed peptide substrate environment. These limitations motivated use of yeast surface display-based directed evolution (3) to create a human ACE2 double mutant (T371L/Y510Ile), referred to as ACE2dm, that is highly specific toward Ang-II relative to apelins and other wild type ACE2 substrates.

Within the collection of cancer models (1,2) in which losartan has been evaluated as a TME-modulating adjuvant that can increase the efficacy of other therapeutic modalities, GBM (2) is a desirable initial venue for demonstrating ACE2dm effectiveness because angiotensin peptides do not readily cross the blood brain barrier (BBB). As such, the brain is a relatively closed compartment within which ACE2dm’s hydrolytic activity can increase the Ang-(1-7):Ang-II ratio local to tumors with reduced influence of mass transfer-associated effects relative those expected for systemic organs in which blood circulation through the vasculature creates angiotensin peptide mass transfer gradients. Such gradients could reduce ACE2dm’s ability to increase tumor-local Ang-(1-7):Ang-II ratio by continuously replenishing Ang-II and shuttling Ang-(1-7) away from the tumor.

To enable systemically-injected ACE2dm to enter the brain we developed a fusion protein, ACE2dm-s1 (Fig 1b), comprised of ACE2dm with a C-terminal (Gly_4_Ser)_4_ linker connecting ACE2dm to the N-terminus of a rat-derived single-chain antibody (scFv), known as 8D3, that binds mouse transferrin receptor (TrfR) and has been used to transport enzymes or other cargo proteins across the BBB in mouse experiments (5). Transient expression in human embryonic kidney (HEK) cell shake flask culture followed by two-stage NiNTA and size exclusion chromatography purification yielded > 25 mg/L of > 90% pure ACE2dm-s1 (Fig 1c). To verify ACE2dm-s1 ability to bind murine TrfR, biolayer interferometry BLI assays were performed (Fig 1d and Supplemental Fig 1). For both immobilized ACE2dm-s1 and TrfR configurations, equilibrium binding dissociation constants (K_D_s) determined by data fitting were single-digit nanomolar; these values align with results for other 8D3scFv-containing fusion proteins (4). To further confirm ACE2dm-s1’s suitability for in vivo studies we used a liquid chromatography assay (3) to measure Ang-II and Apelin-13 hydrolysis rates in vitro and found them to be nearly identical for ACE2dm-s1 and the standalone ACE2dm enzyme domain (Supplemental Figure 2).

To assess ACE2dm-s1’s ability to modify the TME, with a focus on changes in immune cell populations within the brain, we intraperitoneally-injected CT-2A GBM model mice once daily, where this frequency is based on the approximate 10-hour lifetime (5) of 8D3scFv fusion proteins in brain after BBB traversal, with ACE2dm-s1 or PBS negative control vehicle for five days. Subsequently, we conducted flow cytometric analysis of single cells obtained by dissociating brain tissue collected four hours after final injections. CD11b^+^/CD86^+^ and CD11b^+^/CD206^+^ macrophage populations were significantly decreased for ACE2dm-s1 mice (Fig 1e) while CD8^+^/IFN-γ^+^ T cell counts increased approximately fivefold; this increase aligns with CD11b^+^/CD206^+^ macrophage suppressive control over CD8^+^/IFN-γ^+^ T cell populations.

Unpublished results of other mouse studies we have performed showed that, in contrast to ACE2dm-s1, post-insult intraperitoneal injection of standalone ACE2dm had limited or no effect on brain vascular function, brain vascular architecture assessed by immunohistochemistry, and behavioral test readouts in mouse models of stroke and traumatic brain injury (TBI). As such, GBM model mouse studies with standalone ACE2dm were not pursued along with the ACE2dm-s1 experiments described here.

Suppression of activation of stimulator of interferon gamma (STING) signaling is among the mechanisms by which endogenous anti-tumor immune responses can be attenuated within the TME (6). As such, we viewed the STING agonist 2’,3’-cGAMP as an appropriate co-administered agent for establishing ACE2dm-s1’s utility as a TME-modulating adjuvant component of combination therapy regimens that can improve GBM mouse survival. Whereas neither 2’,3’-cGAMP nor ACE2dm-Fc monotherapy significantly increased survival probability relative to untreated mice, survival was improved for the combination therapy group with two of eight mice being viable at the day 50 endpoint (Fig 1f). These results are highly notable given that 100 % mortality was observed for both anti-PD-1 monotherapy and anti-PD-1/losartan combination therapy in this same CT-2A model.

Collectively, our flow cytometry and survival study results bring ACE2dm, and particularly brain-targeted fusion protein versions of the enzyme, to light as a TME-modulating adjuvant with substantial potential as an element of combination GBM therapy. These findings also motivate further work aimed at better understanding the mechanisms of ACE2dm-antibody fusion protein brain delivery, where we recognize that this initial demonstration of ACE2dm-s1 efficacy in GBM mice did not speak directly to whether ACE2dm-s1 entered the brain via TrfR-mediated transcytosis, passive transport, or a combination thereof, and assessing ability to augment ACE2dm migration from brain to blood by fusing ACE2dm to brain-targeting antibodies other than the 8D3 scFv. Additionally, these findings motivate pursuit of ACE2dm, and in particular long-lived systemically-circulating IgG fragment crystallizable (Fc) domain fusion proteins, as TME-modulating adjuvants in combination therapy of systemic solid tumor cancers.

## Supporting information

Supplemental File One

Uncropped Gel Image for Figure 1c

Supporting Data Values

