## Supplemental File One for "Glioblastoma model mouse rescue from mortality by brain-targeted catalytically-enhanced angiotensin-converting enzyme 2 mutant-antibody fusion protein"

**Methods and Supplemental Figures**

Sex As Biological Variable

Sex was not considered as a biological variable for the purposes of this study. The study examined only one sex, i.e., males due to resource constraints. Further experiments would be required to determine the relevance of our findings to female mice.

ACE2dm-s1 Expression and Purification

For ACE2dm-s1 expression and purification, the T371L/Y510Ile mutations described in (1) were introduced into the wild type ACE2 gene. Overlap extension PCR was used to create a ACE2dm-s1 fusion protein gene in which the C-terminus of ACE2dm is followed by a (G_4_S)_4_ linker that connects the ACE2dm catalytic domain to the N-terminus of the 8D3 anti-transferrin receptor single-chain antibody as described (2). The 8D3 scFv gene was obtained via DNA synthesis (Integrated DNA Technologies). The resultant fusion protein gene was inserted between the NheI and BamHI sites of the mammalian secretion vector (Addgene ID 149268) described in (3).

Expi293 cells (LifeTech) were transiently transfected with endotoxin-free DNA (1 mg DNA per liter culture volume) and transfection enhancer added twenty hours after DNA addition. Five days after transfection the cells were centrifuged at 500g for 10 minutes and supernatant was sterilized by sequential vacuum filtration through 0.8, 0.45 and 0.2 µM polyethersulfone (PES) membranes. Affinity purification was performed using ProBond NiNTA resin (ThermoFisher) subsequent to concentration and buffer exchange of NiNTA resin eluate using 30kDa MWCO VivaSpin centrifugal units (Sartorius) and second stage purification with VivaPure MiniH cation exchange columns (Sartorius). Post-ion exchanged protein was desalted into endotoxin-free PBS (pH 7.4) containing 3 µM ZnCL2 to preserve ACE2 activity, sterile filtered with a 0.2 µm polyethersulfone syringe unit, aliquoted, and frozen at -80 C in low protein-binding tubes.

Protein concentration and purity were assessed by SDS-PAGE analysis with coomassie blue staining and GelQuant software (BiochemLab Solutions) was used to quantify band intensities relative to bovine serum albumin standard. Concentrations of final purified ACE2dm-s1 samples are typically in the 3-5 mg/mL range. Purified standalone ACE2dm and wild type ACE2 catalytic domains (1) used for in vitro activity assays were quantified using the methods described above.

Biolayer Interferometry

A ForteBio Octet RED96 system was used to conduct biolayer interferometry assays for quantification of the equilibrium binding dissociation constant (K_D_) for the interaction between ACE2dm-s1 and recombinant mouse transferrin receptor (ACROBiosystems). Assays were conducted in two configurations: ACE2dm-s1 free in solution with TrfR immobilized on the sensor tip and the reverse case. For both configurations amine chemistry was used for protein immobilization with ligands being coupled to sensor tips via incubation in pH 4.0 sodium acetate buffer at a concentration of 10 µg/mL. Analyte loading times of 10 minutes were used with subsequent monitoring of binding signal during a 10-minute dissociation interval. Analyte concentration ranges were chosen based on amount of analyte protein available and an anticipated K_D_ value in the single digit nanomolar range. Binding signal data was analyzed with Octet Analysis Studio software to estimate K_D_ values. Estimated errors in K_D_ values were less than 1 % both assay configurations.

Enzyme Activity Assays

Activity of ACE2dm-s1, standalone ACE2dm catalytic domain, and standalone wild type ACE2 catalytic domain, with the latter two proteins prepared as described (1), was determined by incubating enzymes at a concentration of 700 pM (for Ang-II (Anaspec)) or 1.4 µM (for Apelin-13 (Bachem)) with either 1 µM or 3 µM Ang-II or apelin-13 for 25 minutes for Ang-II reactions or 4.5 minutes for Apelin-13 reactions. Product formation was quantified via liquid chromatography following published procedures (1). Single data points were obtained for each reaction.

CT-2A Glioblastoma Mouse Model

For CT-2A mouse experiments, six- to eight-week-old female C57BL/6J mice (Harlan) were maintained in individually ventilated cages. Animal procedures were performed in accordance with the local guidelines and approved by the Stanford IACUC committee. The immune-competent intracranial CT-2A glioma models were established as previously described (4). In short, mice were stereotactically injected with 5 × 10^4^ CT-2A cells in the right hemisphere with the following coordinates: 3 mm deep, 2.2 mm lateral, and 0.5 mm anterior of bregma. Mice were randomly assigned to treatment groups. On day 7 after tumor implantation, mice were treated with ACE2dm-s1 and/or STING agonist as described in the sections below.

Processing and Flow Cytometric Analysis of Brain Lymphocyte Populations

Beginning on day 7 after tumor implantation, mice (five per group) received intraperitoneal injections of ACE2dm-s1 (3.5 mg/kg) or PBS vehicle. Injections were made once daily for five days and all mice were euthanized four hours after the final injection on day five. Brains were collected and single cells from tumor-bearing whole brains were obtained by mincing tissue via mechanical dissociation using a syringe plunger and 70 µm strainer in RPMI 1640 media (Gibco) on ice. Cells were centrifuged at 1250 rpm for 5 minutes, resuspended in 5 mL of 60% Percoll (GE Healthcare Life Sciences), overlaid onto a two-layer 30% / 70% Percoll gradient and immune cells were collected from the 30% / 70% interphase after centrifuging at 2,000 rpm for 30 minutes at room temperature with deceleration set to zero. Immune cells were next stained with relevant antibodies for flow cytometry (Supplemental Table 1) and cytometry data were acquired using a Cytek spectral flow cytometer with SpectroFlo software (Cytek Biosciences) and analyzed using FlowJo software (BD Biosciences). Gating was performed as follows: [1] exclusion of debris based on forward- and side-scatter characteristics (FSC/SSC), [2] exclusion of doublets using forward-scatter height and width (FSC-H/FSC-W), [3] exclusion of nonviable cells using a fixable viability dye (noted in Supplemental Table 1), [4] identification of the lymphocyte population based on FSC/SSC and CD45^+^ labeling, [5] identification of CD3^+^ T cells and subsequent delineation of CD4^+^ and CD8^+^ T-cell subsets, and [6] quantification of co-signaling receptor expression using gates established with fluorescence-minus-one controls.

**Supplemental Table 1.** Antibody panel for ACE2dm-s1 CT-2A mouse flow cytometry study.

| **Marker** | **Fluorophore** |
| --- | --- |
| Live/Dead | BV510 |
| CD45 | APC-Cy7 |
| CD3 | BV785 |
| CD4 | BV650 |
| CD11b | FITC |
| CD8 | BV711 |
| CD86 | PE Dazzle 594 |
| CD206 | PE-Cy7 |
| IFN-γ | Pacific Blue |
| Foxp3 | Alexa Fluor 647 |

ACE2dm-s1 Combination Therapy Survival Study

CT-2A mice were implanted with tumors as above for the flow cytometry study and assigned into groups of eight mice except for the ACE2dm-s1 only treatment group which contained seven mice. On day 7 after tumor implantation, mice received i.p. injections of ACE2dm-s1 as above or with or without STING agonist (2’,3’-cGAMP (MedChem Express)) administered by intrathecal injection. STING agonist, dissolved in PBS at a concentration of 0.6 mg/mL, was delivered in an injection volume of 5 μL using a 30-gauge needle. Mice were injected with ACE2dm-s1, STING agonist, and/or PBS vehicle as appropriate for the combination and monotherapy groups once daily for five days. Following the fifth day of treatment, survival was determined by daily observation through the study endpoint at day 50.

Statistical Analysis

For flow cytometry studies, comparison of immune subsets between ACE2dm-s1 and PBS vehicle negative control groups was performed with the Student’s t-test and data are presented as mean +/- standard deviation. Survival data was analyzed with the log-rank Mantel-Cox test. Statistical analyses were performed using GraphPad Prism GraphPad Prism version 9.3.1.

SDS-PAGE Image

Uncropped SDS-PAGE gel image for Fig 1c is provided as a separate file.

Study Approval

All mouse experiments were reviewed and approved by the Stanford University IACUC committee and carried out under protocol number 33910.

Data Availability

Absorbance values for the liquid chromatography assay used to measure enzyme activity are reported in the Supporting Data Values file. This file also contains the response readout values for biolayer interferometry assays. Mean fluorescence values for flow cytometry analysis of brain immune cells are also presented in the Supporting Data Value file. Additional data that support the findings of this study are available from the corresponding author on reasonable request.

Acknowledgements

This work is the result of NIH funding, in whole or in part, and is subject to the NIH Public Access Policy. Through acceptance of this federal funding, the NIH has been given a right to make the work publicly available in PubMed Central. PAR was supported by grant R35GM119854. ML was supported by grant R01NS121404.

Author contribution

Conceptualization: PH, ML

Investigation: PH, JC, GRN, JL, SYL

Formal Analysis: PH, JC, GRN

Resources: PAR, ML

Data Curation: PH, JC, GRN

Project Administration: PAR, ML

Funding Acquisition: PAR, ML

Writing - Original Draft: PH

Writing - Review & Editing: PH, ML, PAR, JC, GRN

Supplemental Figures

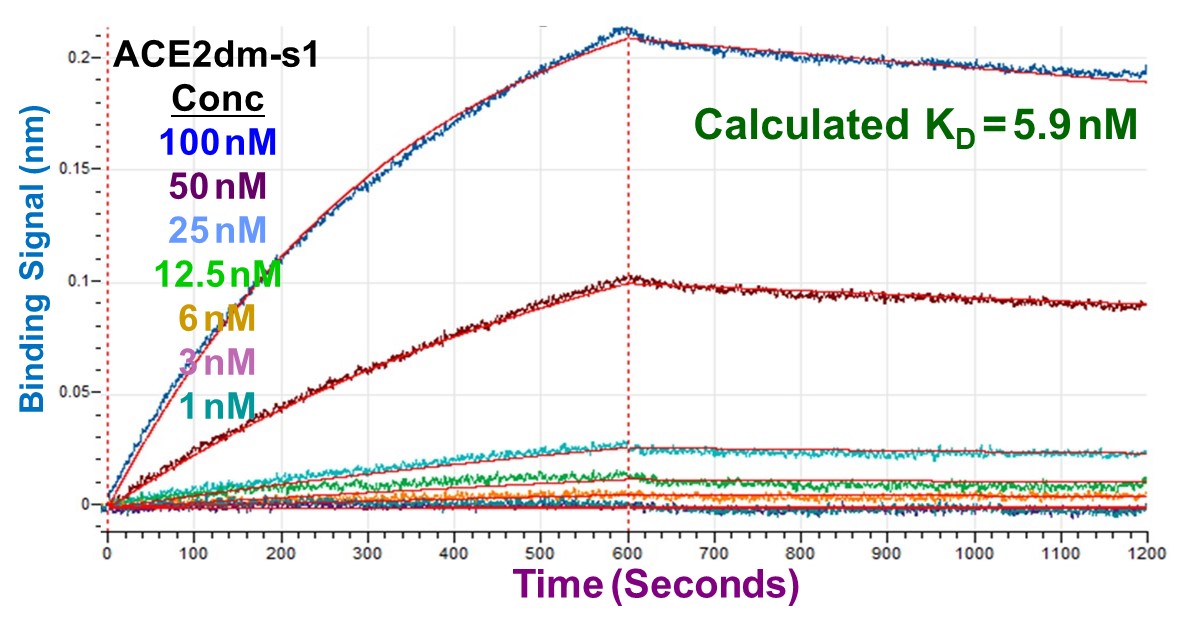

**Supplemental Fig 1.** Biolayer interferometry response curves for mouse transferrin receptor as ligand and various concentrations of ACE2dm-s1 free in solution. Estimated K_D_ value (error was less than 1%) denoted on plot.

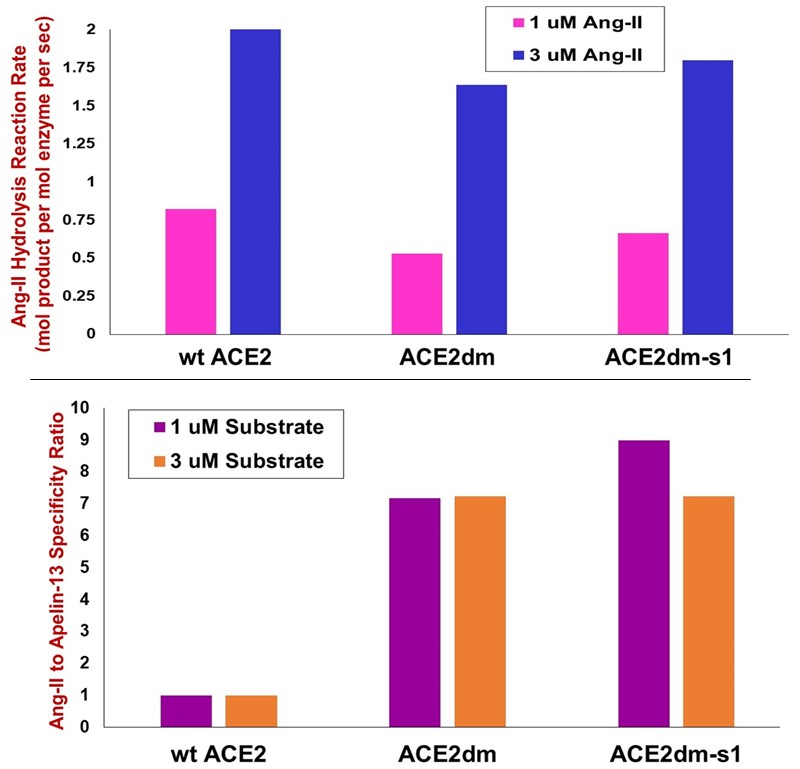

**Supplemental Fig 2.** Upper Panel: Ang-II hydrolysis rates (moles product / moles enzyme / sec) for ACE2 enzymes in reaction buffer at concentration of 700 pM. Lower Panel: Ratios of Ang-II hydrolysis rate to Apelin-13 hydrolysis rate for same ACE2 enzymes assayed in upper panel. Bars represent values for single trials.
