## Supplementary material for "Glioblastoma model mouse rescue from mortality by brain-targeted catalytically-enhanced angiotensin-converting enzyme 2 mutant-antibody fusion protein": Uncropped Gel Image for Figure 1c

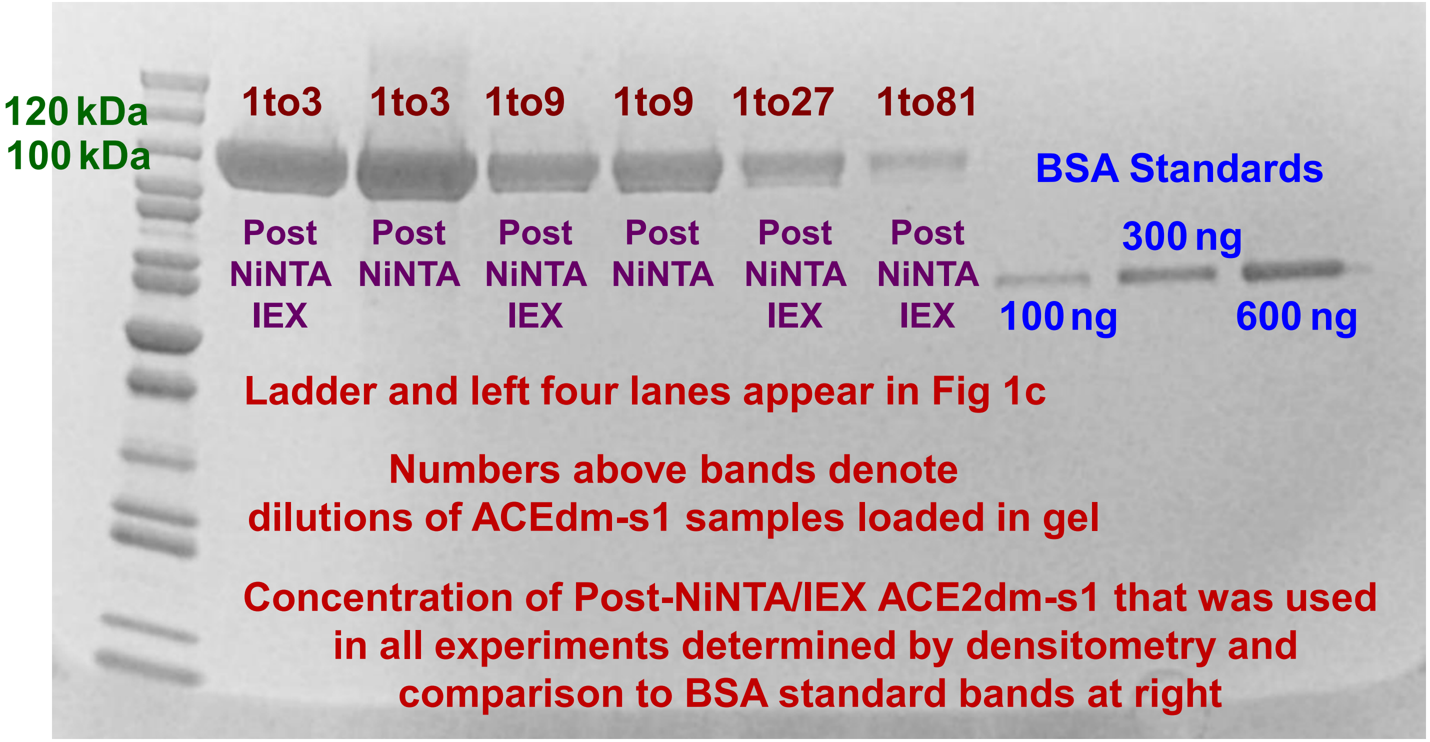
120 kDa
100 kDa
1to3
1to3
1to9
1to9
1to27
1to81
Post NiNTA IEX
Post NiNTA
Post NiNTA IEX
Post NiNTA
Post NiNTA IEX
Post NiNTA IEX
BSA Standards
100 ng
300 ng
600 ng
Ladder and left four lanes appear in Fig 1c
Numbers above bands denote dilutions of ACEdm-s1 samples loaded in gel
Concentration of Post-NiNTA/IEX ACE2dm-s1 that was used in all experiments determined by densitometry and comparison to BSA standard bands at right
